# Historic collections as a tool for assessing the global pollinator crisis

**DOI:** 10.1101/296921

**Authors:** I. Bartomeus, J.R. Stavert, D. Ward, O. Aguado

## Abstract

There is increasing concern about the decline of pollinators worldwide. However, despite reports that pollinator declines are widespread, data are scarce and often geographically and taxonomically biased. These biases limit robust inference about any potential pollinator crisis. Non-structured and opportunistic historical specimen collection data provide the only source of historical information which can serve as a baseline for identifying pollinator declines. Specimens historically collected and preserved in museums not only provide information on where and when species were collected, but also contain other ecological information such as species interactions and morphological traits. Here, we provide a synthesis of how researchers have used historical data to identify long-term changes in biodiversity, species abundances, morphology and pollination services. Despite recent advances, we show that information on the status and trends of most pollinators is absent, but we highlight opportunities and limitations to progress the assessment of pollinator declines globally. Finally, we demonstrate different approaches to analysing museum collection data using two contrasting case studies from distinct geographical regions (New Zealand and Spain) for which long-term pollinator declines have never been assessed. There is immense potential for museum specimens to play a central role in assessing the extent of the global pollination crisis.

## Introduction

Animal pollinators are a critical component of both natural and agricultural ecosystems worldwide, given their role in plant reproduction [1] and food security [2]. As with many other taxa, pollinators are vulnerable to a range of anthropogenic disturbances, which can cause local and regional population declines or even extinctions. The vulnerability of pollinators was identified several decades ago, and was popularized in 1996 by the influential book “The forgotten pollinators” [3]. However, early accounts of pollinator declines were somewhat anecdotal, given the lack of pollinator population data at that time. These initial claims triggered the first efforts to assess this potential issue and included the formation of a US National Academy of Science (NAS) panel in 2006, which was commissioned to assess the extent of pollinator declines. The NAS report concluded that “For most pollinator species […] the paucity of long-term population data and the incomplete knowledge of even basic taxonomy and ecology make definitive assessment of status exceedingly difficult” [4]. Since then, studies on pollinator responses to various global change drivers have multiplied rapidly. Researchers have now developed strong consensus that disturbances such as habitat destruction, land-use intensification, chemical exposure, exotic species and climate change are causing pollinator declines, and often act synergistically [5,6]. Yet, the current status and population trends of most pollinator species worldwide remain unknown. For example, a recent IUCN report concluded that even for Europe’s comparatively well-studied bee fauna, greater than 55% of bee species fell into the “Data Deficient” category [7]. For countries outside of Europe and the US, data on pollinator populations is almost non-existent.

One of the main barriers to identifying long-term pollinator population trends is that pollinators are incredibly taxonomically diverse and include bees, flies, butterflies, beetles, birds, bats and lizards [8]. Additionally, many pollinators are highly mobile, short-lived and small, which makes monitoring their populations difficult. Bees are generally regarded as the most important pollinator group due to their abundance, pollination efficiency and widespread distribution [9]. However, bees are diverse, with more than 20,000 species currently described worldwide, and often require expert taxonomists for identification. Furthermore, the uneven distribution of researchers has resulted in geographical biases in bee decline research [10], as well as taxonomic biases toward species that are easier to identify, such as bumblebees [11,12].

One solution to overcoming these barriers is the use of space-for-time substitutions, where researchers compare pollinator populations across environmental gradients. Despite critiques on the robustness of this approach [13,14], these studies currently provide the most extensive source of pollinator population data. For example, researchers have recently estimated bee richness declines for every country in Europe using predictions from models of pollinator associations with different land-use types [15]. A second important method is the use of data collected from pollinator monitoring programs, which are often driven by citizen scientists. This approach was inspired by successful butterfly monitoring programs [16] and is currently being extended to other pollinator taxa. However, these programs require significant time to generate long-term datasets and cannot be used to assess historic pollinator populations. Finally, the most practical approach for assessing long-term historical pollinator population trends is to use historical information on species occurrences, which is often archived in museum collections [e.g. 17].

In this review, we first assess current evidence for pollinator richness declines and present a roadmap outlining a strategy for using historical collection data to fill current knowledge gaps. We highlight the major technical difficulties involved in using historical collection data and demonstrate several approaches for analysing different types of collection data to assess long-term pollinator population trends. Finally, we highlight the need to move beyond simple biological diversity descriptors and unleash the power of historical data to assess changes in species interactions, ecosystem functioning and evolutionary changes through time.

## Current evidence on pollinator declines

At a global scale, current evidence of pollinator declines is highly limited with most data restricted to the US and Europe. It is unsurprising that studies on pollinator declines are biased towards developed western countries, which have also been subject to extensive anthropogenic disturbance. For example, in the UK and the Netherlands, a citizen science based study using both observations and museum collection data detected strong richness declines for bees, hoverflies and flowering plants [18]. In the Netherlands, museum data have also revealed simultaneous plant and pollinator declines [19]. Specifically, bee species with the strongest host plant preferences (i.e., specialists) displayed the strongest declines and thus, were most threatened with extinction. However, it is important to note that even for these two countries, local estimates of pollinator richness are biased toward large cities and regions dominated by agriculture, and thus lack data for well-preserved natural areas. Further exploration of this dataset revealed that for declining pollinator taxa, the trend has attenuated in recent decades [20].

Although studies of local pollinator communities often detect richness declines, regional richness may remain relatively stable. For example, regional estimates for bee species richness changes in the eastern US show moderate declines [17] and very few regional extinctions [21]. This is a pattern also detected in the UK, where relatively few regional bee extinctions have been reported [22]. These regional findings are in stark contrast with the widespread local extinctions reported in local studies. For example, Burkle et al. [23] compared historical observations of bee species’ occurrences in a large forested ecosystem with remaining forest remnants and reports several local extinctions. However, it is important to note that there is strong concordance between local extinctions and regional declines [24], suggesting that local extinctions are indicators of regional population declines.

Reported declines for bumblebees are the most severe of all pollinator taxa. For example, declines of up to 18% in local bumblebee richness have been reported for Belgium and the Netherlands [20]. In other parts of Europe, local richness declines range from 5% in Great Britain [20] to 42% in Denmark [25]. In the USA, reported bumble decline are also severe with estimates ranging between 25% [26] and 30% [17]. However, studies on species richness changes for other pollinator taxa are both scarce and geographically restricted. For butterflies, the only evidence of richness declines comes from Europe. Butterfly species richness has declined substantially in the Netherlands and Belgium since the 1950’s, although declines in Great Britain have been less severe [20]. In Belgium, another study [27] found that richness declines have been severe (approximately 30%), although this study assessed richness changes over a longer time period (early 1900’s to 2000) compared to [20] (1950-69 vs. 1970-80 and 1970-89 vs. 1990-2009). In parts of Germany, up to 70% declines in local butterfly richness have been reported [28]. Compared with other insect pollinator taxa, there are very few studies on hoverfly species richness changes, which are all restricted to Europe. In Belgium, Great Britain and the Netherlands, hoverfly richness changes have been modest [20]. In the Netherlands, moderate increases in hoverfly species richness have been shown, whereas in Great Britain no significant directional changes were detected [18]. Furthermore, directionality (richness increase or decrease) varies depending on the time period assessed. For example, hoverfly richness decreased in Belgium by approximately 6% from 1950-69 to 1970-80, but increased by approximately 10% between 1970-89 and 1990-2009 [20].

For illustrative purposes, we mapped the findings of this studies in Figure 1 to show the strong contrast between bee species richness worldwide, with bee diversity hotspots in Mediterranean countries, against the paucity of countries for which we have any local or regional data on bee, hoverfly or butterfly declines (see raw data in Sup Mat 1). Despite outside of Europe and the US and for non-insect taxa, there are very few or no studies on pollinator declines using historical records, there are species-specific examples of historical losses from different parts of the world (e.g., *Bombus dalbhomi*; [29]).

**Figure 1:**
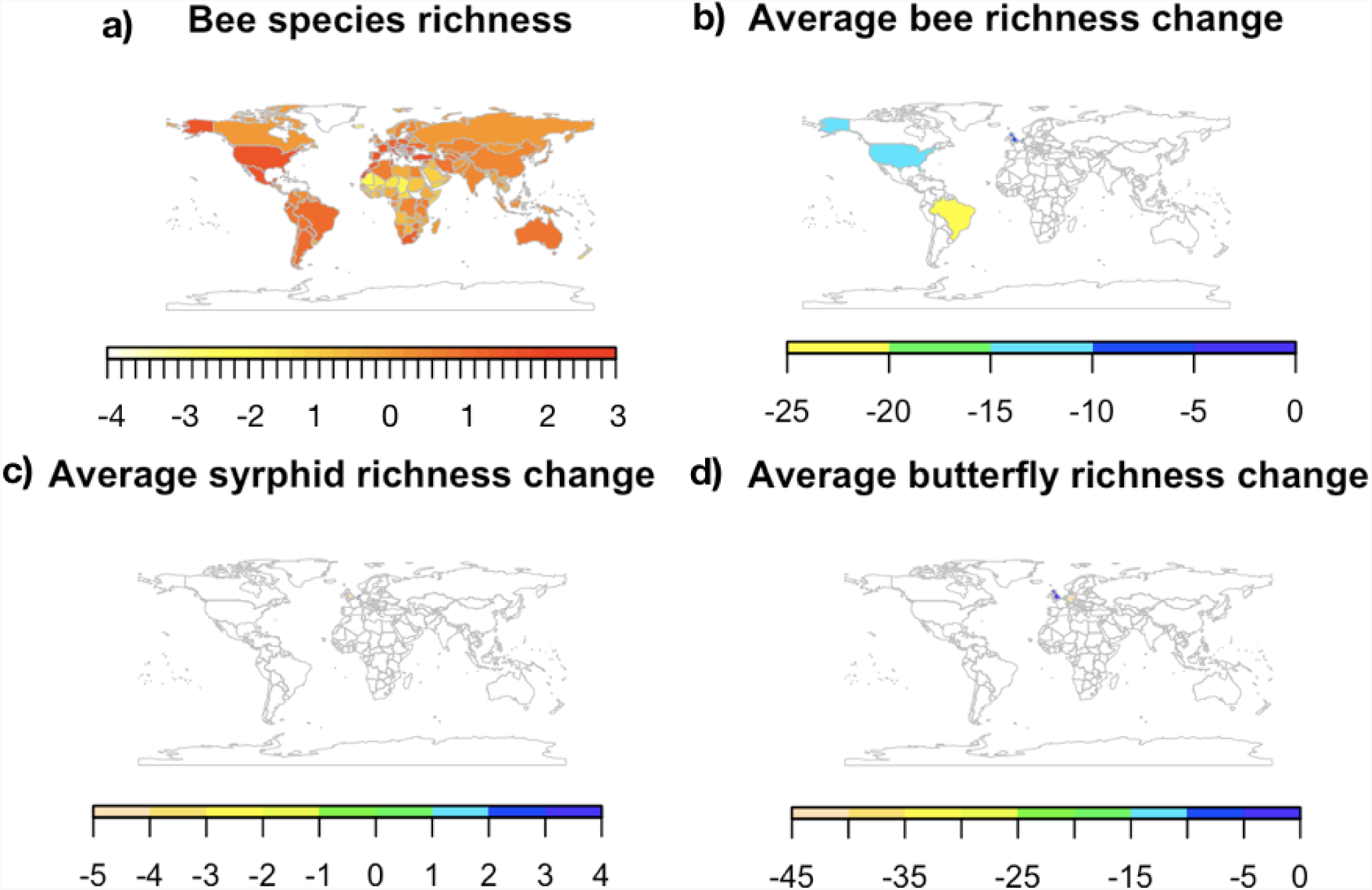
Global map showing a) bee species richness per area (Data from www.discoverlife.org) calculated as the residuals of the log-log regression between bee species richness per country and country size. This correction accounts for the species-area relationship. Warmer colours indicate higher bee diversity. Note that some African countries may have incomplete listed faunas and that Alaska is included with USA values. Countries with available historical changes in (b) bee, (c) hoverfly and (d) butterfly richness within the last 100 years. Warmer colours indicate steeper average declines. Countries without data are coloured in white.

## Using historical collection specimen records to fill knowledge gaps

Estimates of pollinator declines are lacking for most countries worldwide (Figure 1). The use of historic collection data may be the most effective tool for filling these gaps. The core aim of museums is to conserve and curate historic collections. Thus, they serve as a precious repository for specimens, and at the same time, often ensure higher quality taxonomic identification. Yet, the major bottleneck for researchers wanting to use these data is the lack of digitization. Digitizing old collection specimens is not a trivial task and requires expertise to (i) ensure proper taxonomic identification [30–32], (ii) geo-locate the coordinates of collection events (e.g. http://www.geonames.org) and (iii) store the data in a properly curated database [33]. Undertaking this process for tens or hundreds of thousands of museum collection specimens can be a daunting task and requires specialized personnel. While some tasks can only be undertaken by people with specialist skills (e.g., taxonomists), new technologies and citizen science can speed up the collection digitization process. High resolution photos of specimens and associated labels can be uploaded to the internet, where the task of image transcription can be distributed across hundreds or thousands of volunteers (e.g., https://www.zooniverse.org/). In addition, new algorithms have been created that allow location geo-referencing based on vernacular names (e.g. https://geoparser.io). However, achieving this requires adequate funding [34].

Where digitization has been completed, the data provide a rich source of information, allowing assessment of the current status and long-term trends of pollinator populations [17,19,35]. This is despite the fact that museum collections often have a number of biases, including unknown sampling effort, personal interests of collectors and the curatorial techniques used. For example, collectors tend to target rare or unusual over common taxa, discard damaged individuals or only accession a certain number of individuals. In addition, collections are often made opportunistically, leading to a spatial biases where difficult to access areas are under-sampled or conversely, where samples are biased towards easily accessed locations (e.g., towns/cities and/or roadsides). Further, museum collection data can only be used to determine where species are present and not where they are absent. However, given adequate sample sizes and appropriate statistical techniques, most biases can be accounted for [e.g. 17,36,37].

## The way forward: Prioritizing the low hanging fruit

As we have shown, there is a paucity of countries for which historical data is available (Figure 1), and hence can be used as baseline for assessing pollinator population declines. While ideally one would aim to digitize all museum collection records, this is unlikely in the near future, predominantly due to funding constraints. Here we show how researchers can optimize the use of historical collection data to assess long-term pollinator population changes.

GBIF (https://www.gbif.org/) is a central repository for global species occurrence data. Much of these data come from museums, private collections and government research institutes, but several other sources are also integrated. In combination with the popular statistical language R [38], GBIF can be directly queried into your computer [39] and data availability can be checked for the region of interest. Focusing on bee taxa, we show here the number of modern and historic bee records currently available for different countries (Figure 2a, Sup mat 2). Thirty-seven countries have more than 1800 records in each time period, making these data potentially analyzable without further data collection effort (see Figure 2b and c for an initial exploration). However, a proper analysis of this dataset would require a careful inspection of the data, as we detail below for two specific countries (Spain and New Zealand). In contrast, most countries fall short in one or both axes of Figure 2a. For example, a variety of countries located in different continents such as Switzerland, Sri Lanka, Nicaragua or Zimbabwe have a decent number of recent records, but lack historical collections. In this cases, researchers should prioritize the digitalization of old material before embarking on data analyses. For this end, it is also important to note that historical records are not always vouchered in local museums (i.e., many European and USA museums contain large collections of pollinators from other countries). Finally, it’s remarkable that more than 192 countries have less than 1000 records for each of both time periods, making them poor candidates for analysing long-term pollinator population trends. Aside from bees, similar exploratory analyses can easily be conducted for other taxa.

**Figure 2.**
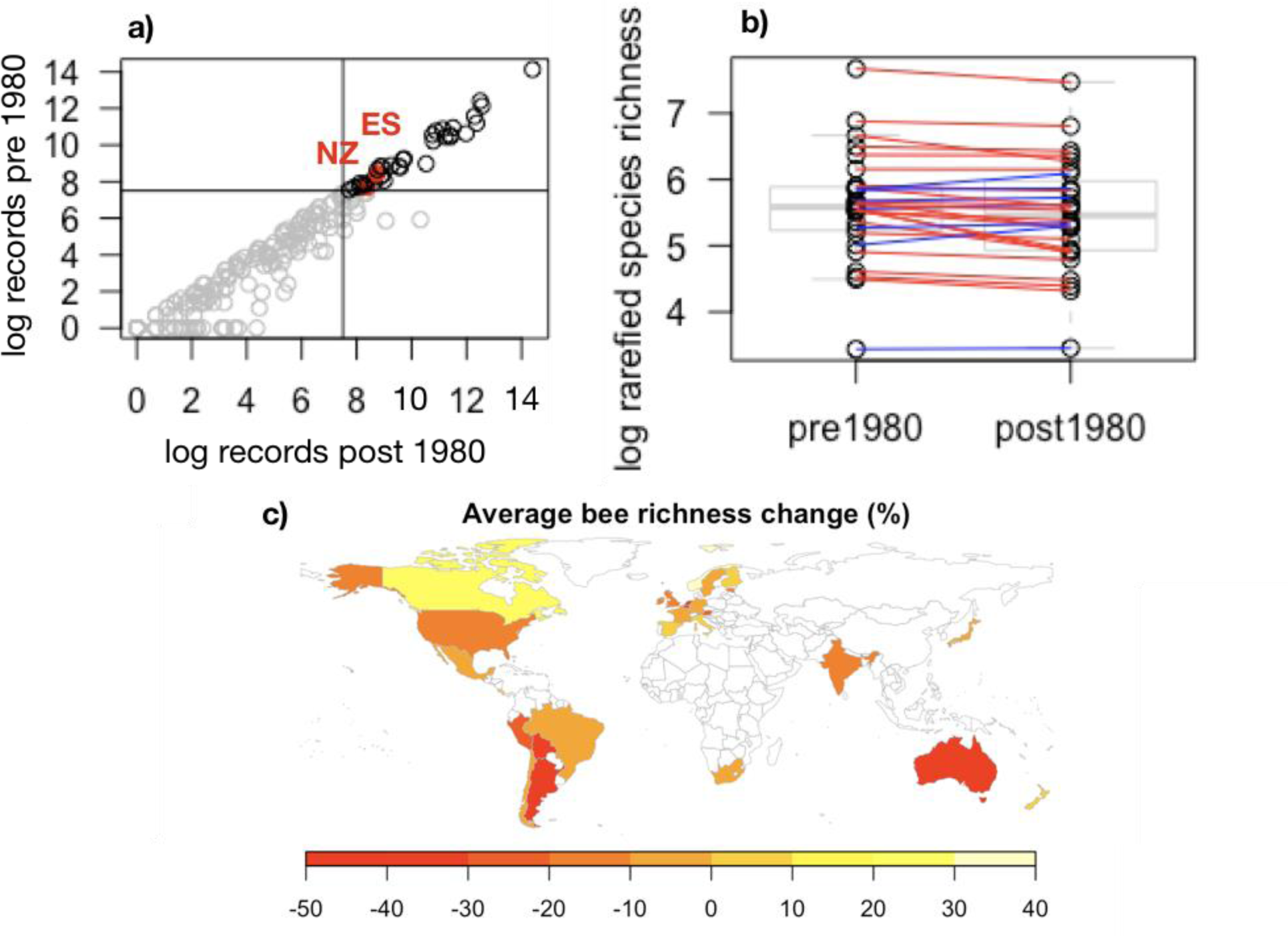
Exploration of available data for bee records showing: (a) The number of bee occurrences before 1980 and after 1980 in GBIF for each country. The upper right quadrat (records in black) contains well covered countries with New Zealand (NZ) and Spain (ES) marked in red (see below). For well covered countries, we show preliminary comparisons of the rarefied number of species in both time periods and show that for most countries (21 out of 28) the number of species recorded is slightly lower (average of 10% richness decline; red lines) for recent time periods (b). Data is log transformed for visualization purposes. A more careful analysis of this data would help complete the map of global declines (c). In this map we plot the % change in species recorded in GFIF for the available countries to show the potential geographic coverage. Note that this data is likely to contain strong undetected biases, as we explore below.

As stated above, once historical collection datasets are made available, researchers must identify any potential biases. We explore this process with two contrasting dataset examples (Spain and New Zealand). In the Spanish dataset, most of the data comes from a few specific locations and was collected by a few specific teams. Hence, the geographical coverage is not representative. Even worst, historical and modern collections do not overlap spatially, making any inference impossible to interpret. In this case, we contacted the original collectors of the historical data to define their sampling protocols. We then resurveyed the same sites (35 years after the original surveys) using the same sampling protocols. In contrast, the New Zealand dataset includes a wide suite of collectors and collection locations but shows no obvious biases in geographical and taxonomic coverage through time. We complemented GBIF data with further museum collections for bees and flies and analyze the regional richness changes through time. For these two case studies, we provide annotated R scripts as examples of analysis for different dataset types (Sup Mat 3). These different analytical approaches allow us to reveal long-term trends in pollinator populations for regions with contrasting sampling histories. We hope this resource will encourage researchers to analyse data for regions where current information on pollinator declines is lacking.

## Case study one: Spain

Spain provides an interesting study system because its natural habitats have been transformed extensively by humans over a long time period, but land-use is not as intensive compared with many other European countries. In addition, Spain is a bee diversity hotspot (Figure 1a) and maintains a relatively heterogeneous landscape. Spain has already digitalized a large amount of pollinator occurrence data for both historic and recent periods (Figure 2a). However, visual inspection of the data revealed clustering around a few localities. Further, historic records did not spatially match recent records, making comparisons difficult. For this dataset, most of the historic records were located around Valladolid and were collected by Enrique Asensio and collaborators. There has been no recent sampling of bees in this area. However, we found that Enrique systematically sampled six independent locations and that additional historical data were available at the “Museo Nacional de Ciencias Naturales” and other minor collections. Digitization of these records, along with a re-survey of the original sampling locations provided an excellent dataset for a before and after comparison of bee communities.

In brief, after cleaning taxonomic names for possible typos and synonyms using the taxize package [40], we checked for sampling completeness for both time periods and compared rarefied species richness for each site before and after 1980 with a paired t-test (rarefication at 1000 specimens). We found that there were a reduced number of species at sites after 1980 (mean difference 20.27 species; 95 percent confidence interval: -1.03, 41.58; t = 2.44, df = 5, P = 0.06). However, this trend was highly dependent on site identity, as two out of six sites showed no richness declines. Interestingly, these two localities were the two that has experienced less land use changes (both are natural areas embedded into agro-ecosystems). In contrast the other 4 localities suffered large urban or agricultural intensification. In addition, species lost in the re-surveys are not a random selection of species, but are clustered in a few genera. For example, Andrenidae and their parasites (e.g. *Nomada*) showed the strongest declines whereas Halictidae tend to be more stable (Sup mat 4). This pattern of winners and losers of land use intensification is in accordance with findings elsewhere [17], indicating that some clades are more sensitive to disturbance than others.

## Case study two: New Zealand

In contrast to Spain, New Zealand is an isolated oceanic archipelago, with a distinctive pollinator biota and a unique history of human occupation. Much of New Zealand’s pollinator fauna is also relatively depauperate. For example, New Zealand has only 27 native bee species [41], which is a fraction of nearby Australia’s c. 1600 species [42]. However, New Zealand has a surprisingly high diversity of flies (Diptera), which are important pollinators in many ecosystems [43]. Thus, New Zealand provides a unique system to study long-term changes in pollinator communities, and is unlike continental Europe and the US, which have been the focus of an overwhelming majority of pollinator decline studies.

In global terms, human colonisation of New Zealand was relatively recent (c. 740 y) [44]. Before human arrival, New Zealand was predominately forested, but has since been dramatically altered by people. Early Māori settlers cleared forests by burning and more recently, European colonists cleared large tracts of remaining forests and drained low-lying wetlands for agriculture, mostly before 1900 [45]. Therefore, human activity likely affected pollinator communities in New Zealand long before surveys and specimen collections began. Nevertheless, we can use museum records to identify trends in pollinator communities during New Zealand’s more recent history.

We used New Zealand bee collection records gathered from multiple sources, including university, research institute, museum and private collections. Collection records from the New Zealand Arthropod Collection (NZAC) are freely available online (https://scd.landcareresearch.co.nz/). Fly pollinator data was obtained from three participating New Zealand museums and covers two families (Calliphoridae and Syrphidae) that contain important fly pollinators. Collections for the bee and fly datasets span over 100 years (early 1900s to late 2000s).

We followed protocols outlined in [17] to analyse the New Zealand data at the regional level. First, we filtered our original datasets so that data used for analyses only included independent collection events. To do this, we removed specimens collected at the same location, on the same date, and by the same collector. We found our data had reasonable coverage across time periods, although there was a peak in collection occurrences from 1960-1980. Further exploration of the New Zealand native bee data raised doubts on collection completeness in records prior to 1970, so we removed these records from further analyses. We accounted for differences in collection effort through binning collection records by time so that each bin had a similar number of records but a different number of years. We then estimated richness for each time period bin by rarefying all bins to an equal number of specimens and calculated the mean species richness ±SE for each bin. Finally, we estimated the significance of change in richness using a permutation test that randomly reordered time periods and calculated the correlation between time period and species richness. Thus, reported P-values were the proportion of permutations that had higher or lower correlations compared to the correlation between richness and the actual chronological time period sequence.

Second, to determine if the probability of finding a species in the collection changed over time, we used a general linear model with a binomial distribution and a logit link. For species that showed overdispersion, we used a quasi-binomial distribution. Further, we only included species in this analysis for which we had 30 or more records. To account for differences in sampling effort between years, we weighted each year by the total number of samples collected that year.

We found that rarefied richness for native bees was stable through time. Exotic bees showed an increase in rarefied richness, but this trend was non-significant (P-value for both natives and exotic bees > 0.05). In contrast, native fly richness declined, whereas exotic fly richness increased, although results for these groups were also non-significant (P-values for both groups > 0.05). Note that rarefied richness is sensitive to species evenness, so increases in rarefied richness over time may actually indicate increased species evenness and vice-versa for decreased richness.

However, at the species level, we found that 11 out of 27 bee species increased in relative occurrence over time (10 native and one exotic) and three bee species declined in relative occurrence (one native and two exotic) (Figure 3). Interestingly, the two exotic bee species that declined in relative occurrence were both in the genus *Bombus*, which were intentionally introduced into New Zealand for the pollination of crops. Native bees that increased in relative occurrence were mostly from the genus *Leioproctus*, which are medium sized, ground nesting solitary bees. Only one out of 14 fly species increased in relative occurrence, which was exotic, whereas four species decreased in occurrence (three native and one exotic). Native flies that decreased in relative occurrence were all Syrphidae in the genus *Helophilus*.

**Figure 3.**
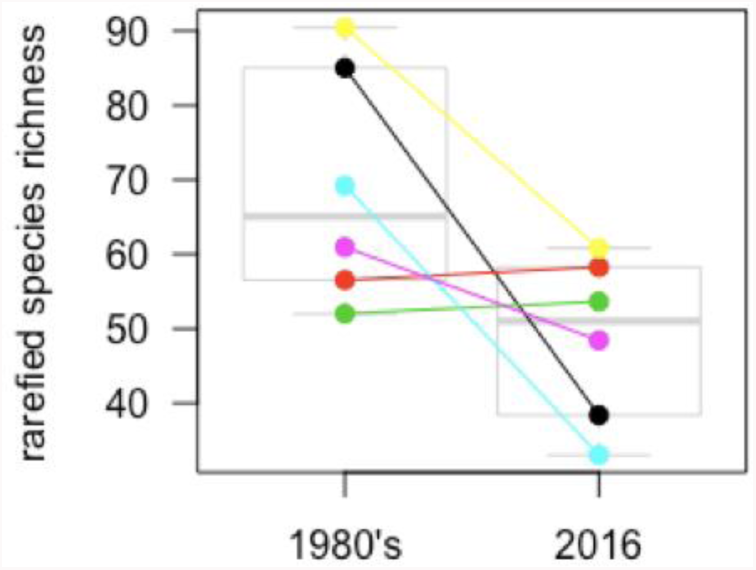
Comparison of historic collections (1980’s) and modern re-surveys (2016) of the rarefied richness of bees at six Spanish localities.

**Figure 4.**
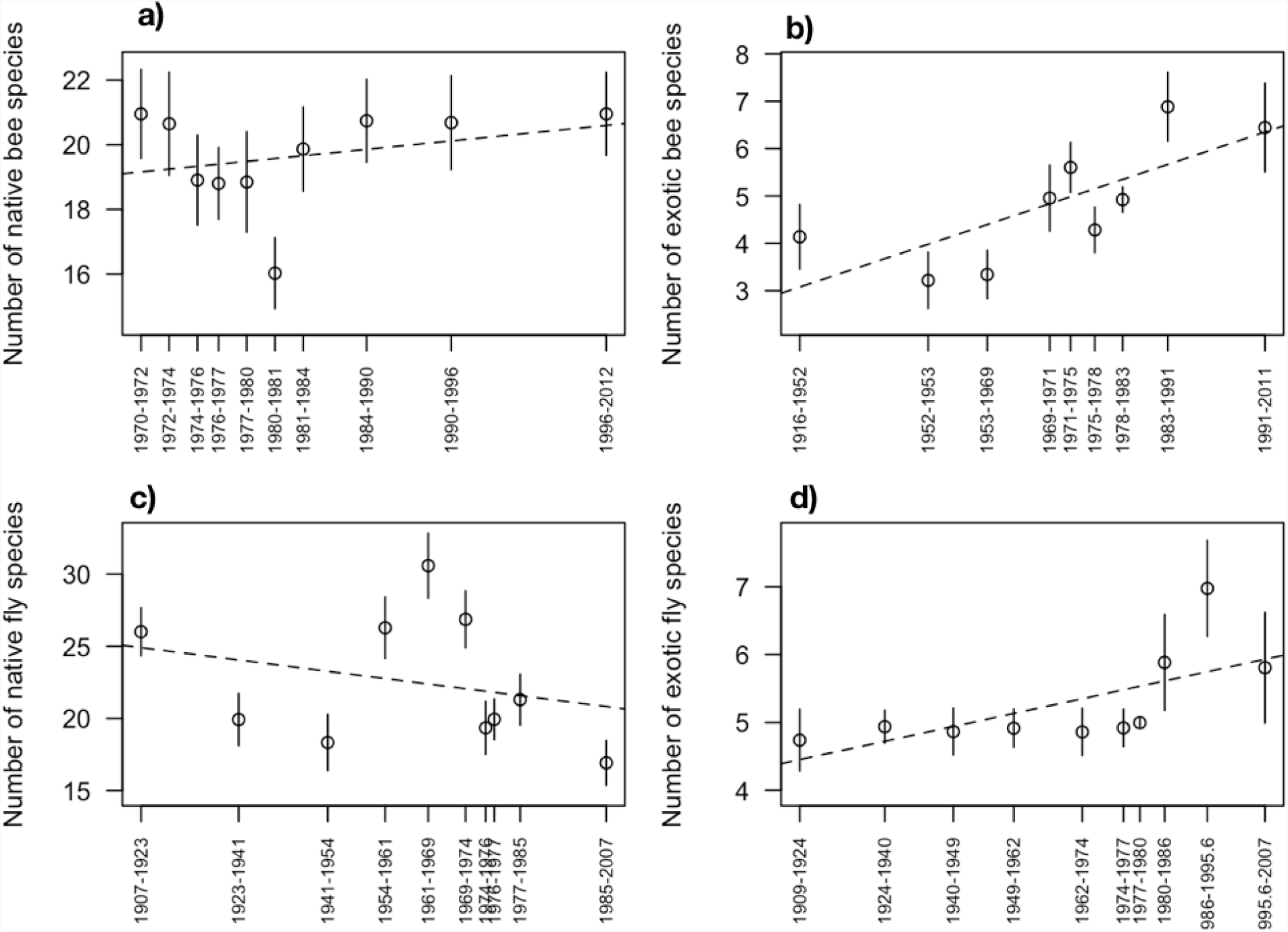
Changes in rarefied species richness for different pollinator groups in New Zealand over time. All trends were non-significant (α = 0.05).

**Figure 5.**
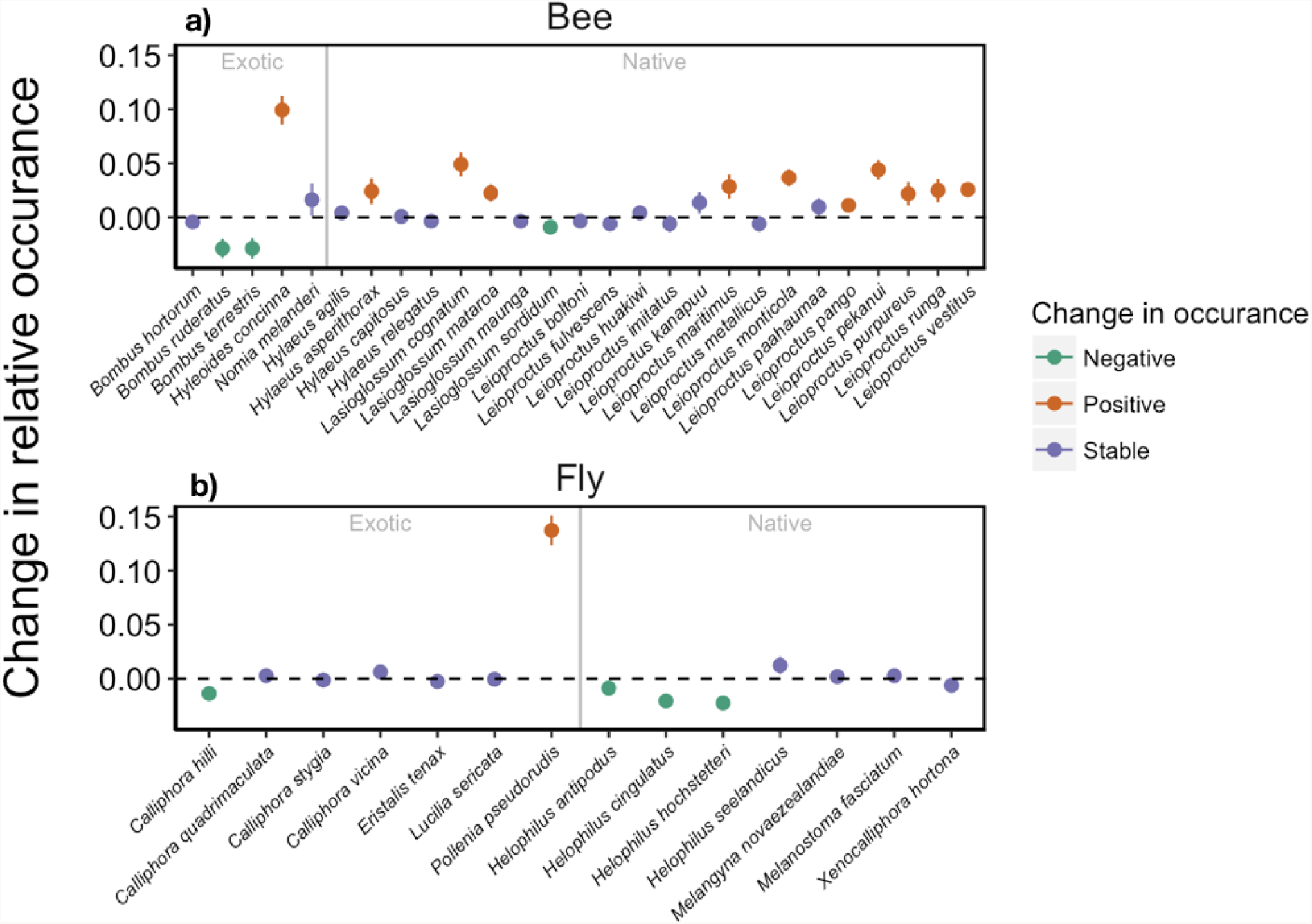
Model estimated changes (± 1 SE) in the relative occurrence frequency of different New Zealand bee and fly species in museum collections over time.

## Beyond species occurrences

A recent study found that more than 90% of the papers investigating pollinator responses to land-use change focused solely on richness and abundance descriptors [9]. But in addition to local (*alpha*) diversity and regional (*gamma*) diversity, researchers need to assess changes in turnover between sites (*beta* diversity). Environmental changes often result in a few “winner” species and many “losers” species [17]. Identifying winners and losers is critical as the few winners are often exotic and represent a subset of traits that facilitate survival in highly modified environments [46]. These changes can have important effects for pollination of native plant species and crops [47].

In addition, museum specimen collections can provide much more information besides species occurrence records, given that such information is recorded when digitizing collections. This is particularly important for identifying mechanisms of decline and adaptation. For example, recording the date of collection is particularly important for tracking of phenological advances congruent with contemporary climate change [48]. In addition, pollinator specimen labels often include information about the host plant on which the specimen was collected. This information critical for understanding past and present species interactions [49]. Aside from this information, bee specimens often contain pollen loads trapped on hairs, from which past visitation events can be identified [50]. Finally, museum specimens can be measured to track evolutionary changes by measuring the traits of specimen. This approach has been already used to investigate tongue length [51] and body size [52] changes in response to climate and land-use change. Finally, plant herbariums can also contain indirect evidence of pollinator and pollination declines [53], a basic information for linking pollinator declines with its consequences for ecosystem functioning.

## Conclusions

Unleashing the power of museum collection data to answer pressing ecological and evolutionary questions is at our hands, but requires the coordinated effort of many actors. Using two case studies, we show that strong collaboration between museum curators and ecologists is key to understanding data and treating it appropriately. To progress our understanding of the global pollination crisis, researchers and curators must aim to digitize museum collection data and make it readily available in a format that is widely accessible. Centralization of regional and national museum collection data in existing global platforms, such as GBIF, would facilitate free and widespread access. However, datasets could also be stored in alternative webpages or database repositories (e.g., university and museum webpages or Dryad) providing they are thoroughly documented and easily retrieved and combined with other datasets using open science tools [54].

We must revolutionize the way that researchers collaborate with museums, in order to foster healthy bidirectional relationships. For example, ecological researchers collect massive amounts of specimens, but these are often inappropriately vouchered [55,56], rendering them less useful for future research. To improve this process, strong communication between museums and researchers is required. However, this can only be achieved with adequate funding and recognition that accurate data recording and long-term preservation are critical for research [57].

To identify global trends in pollinator declines we require robust data, collected from diverse geographic regions. It is also crucial that these data are analysed appropriately. This requires researches to identify biases and to any fill taxonomic and geographic gaps where possible. We need to place increased emphasis on quantifying pollinator declines in regions outside of the US and Europe, and for pollinator groups other than bees. For the US and Europe, there have been few regional bee extinctions [17,22] but in disturbed ecosystems, declines are widespread [15,18]. For most other pollinator taxa and regions throughout the world we know almost nothing. Moving forward, the first step for many taxa will be to identify and describe species. Only then can we begin to document pollinator declines.

## Acknowledgements

We thank Curro Molina, Carola Warner, Patrick McQuinn, and Crona McMonagle for data entry and Gregorio Aguado for carrying out the Spanish re-sampling. We thank Barry Donovan for providing New Zealand bee collection records and E. Asensio for sharing his historical data and knowledge. We thank the “Museo Nacional de Ciencias Naturales”, specially Mercedes Paris, ITACyL (Instituto Tecnológico Agrario de Castilla y León), Canterbury Museum, the New Zealand Arthropod Collection and the Museum of New Zealand Te Papa Tongarewa for access to historical collections. IB was funded by a “Fundación Banco Bilbao Vizcaya Argentaria” (FBBVA) project. DW was funded through Landcare Research within the Characterising New Zealand’s Land Biota Portfolio.

The datasets supporting this article have been uploaded as part of the supplementary material and will be deposited at dryad or Figshare upon acceptance.

We have no competing interests

Author contributions
IB wrote the initial draft. DW and OA provided data. IB and JS analysed the data. All authors contributed to writing the manuscript.

